# High cell diversity and complex peptidergic signalling underlie placozoan behaviour

**DOI:** 10.1101/360925

**Authors:** Frédérique Varoqueaux, Elizabeth A Williams, Susie Grandemange, Luca Truscello, Kai Kamm, Bernd Schierwater, Gáspár Jékely, Dirk Fasshauer

**Author notes:** **Correspondence** (GJ), (DF).

## Abstract

Placozoans, together with sponges, are the only animals devoid of a nervous system and muscles, yet both respond to sensory stimulation in a coordinated manner. How behavioural control in these free-living animals is achieved in the absence of neurons and, more fundamentally, how the first neurons evolved from more primitive communication cells during the rise of animals is not yet understood [1–5]. The placozoan *Trichoplax adhaerens* is a millimeter-wide, flat, free-living marine animal composed of six morphologically identified cell types distributed across a simple bodyplan [6–9]: a flat upper epithelium and a cylindrical lower epithelium interspersed with a loose layer of fiber cells. Its genome encodes several proneuropeptide genes and genes involved in neurosecretion in animals with a nervous system [10–12]. Here we investigate neuropeptide signalling in *Trichoplax adhaerens*. We found specific expression of several neuropeptides in non-overlapping cell populations distributed over the three cell layers, revealing an unsuspected cell-type diversity of *Trichoplax adhaerens*. Using live imaging, we uncovered that treatments with 11 different neuropeptides elicited striking and consistent effects on the animals’ shape, patterns of movement and velocity that we categorized under three main types: (i) crinkling, (ii) turning, and (iii) flattening and churning. Together, the data demonstrate a crucial role for peptidergic signalling in nerveless placozoans and suggest that peptidergic volume signalling may have predated synaptic signalling in the evolution of nervous systems.

## RESULTS AND DISCUSSION

### *Trichoplax adhaerens* comprises multiple populations of peptide-secreting cells

Peptidergic communication is widely used throughout most of the animal kingdom and may even predominate neural communication in phyla such as cnidarians and ctenophores [11,13], but its importance in placozoan physiology and behaviours is not known. The genome of *T. adhaerens* encodes for determinants of both chemical and peptidergic (neuro-)transmission [10–12]. Among the regulatory neuropeptides predicted from the genome [11,12], we selected candidates with good antigenic profiles and generated antibodies against five peptides bearing an amidation site (SIFGamide, SITFamide, YYamide, RWamide and FFNPamide) as well as against one of the four predicted insulin-like peptides, TrIns3. Immunostainings with the purified antibodies in whole-mount *T. adhaerens* uncovered six populations of cells with distinct distribution patterns (Figure 1A–C). Upon co-labelling experiments [14], these peptides were never found to be co-expressed, as illustrated for SIFGamide, SITFamide and FFNPamide in Figure 1B. Instead, it appears that each of the 6 populations of neuropeptide-secreting cells spreads concentrically, delineating distinct territories (Figure 1C). Starting from the outside of the animal toward its inside, SIFGamide immunoreactive cells are found at the very edge and the outer rim of the upper epithelium; in the upper epithelium also, SITFamide cells distribute further inwards in an equally broad annular area. Together, the description of these populations validates and refines the suggested rim-vs-center cell composition of T. *adhaerens* [7,8]. Intriguingly, TrIns3-labeled cells are located in the lower epithelium along a thin ring across the boundaries of SIFGamide and SITFamide territories; a few, small TrIns3-reactive cells are at times also found at the edge.

**Figure 1.**
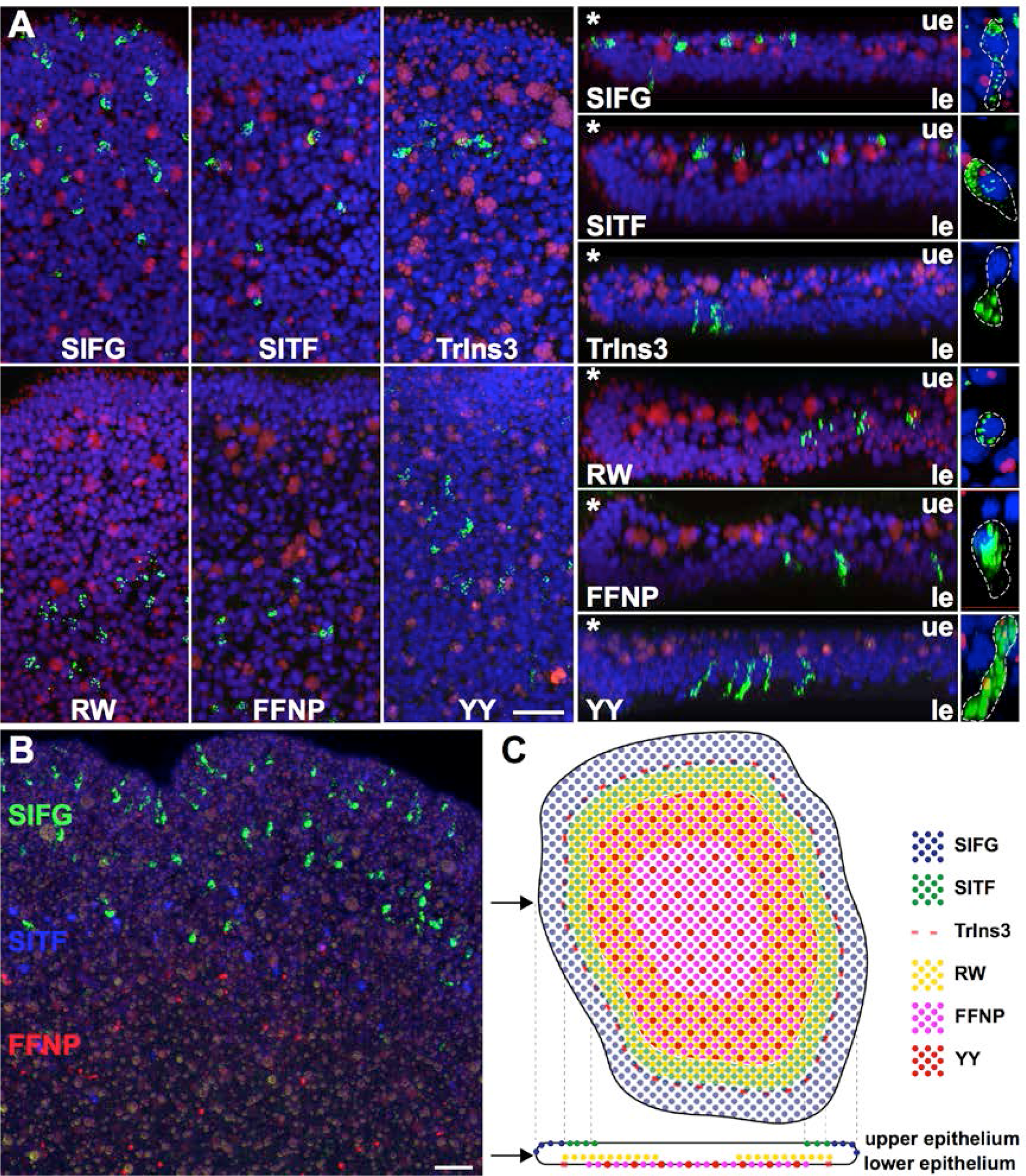
Distribution of peptide-secreting cells in *Trichoplax adhaerens*. (A) Cells expressing SIFGamide, TrIns3, SITFamide, YYamide, RWamide and FFNPamide are spatially organized. Maximal intensity projection images illustrate populations of labelled cells (green) for the indicated peptides upon whole-mount immunostaining of *T. adhaerens* individuals. Images show the cumulative positions of labeled cells from above (left panel) and from the side (in a 10 µm slice; middle panel); representative cells of each population are shown at high magnification in their long axis (horizontal for SIFGamide, SITFamide and RWamide cells, vertical for TrIns3, FFNPamide and YYamide; right panel); labeled granules widely vary in size and distribution across cell populations. Hoechst-stained nuclei (blue) and autofluorescent particles (red) serve as landmarks to locate upper and lower epithelial layers. (B) Populations of peptidergic cells do not overlap. A maximal intensity projection confocal image illustrates the partitioned distribution of SIFGamide (green), SITFamide (blue) and FFNPamide (red)-immunoreactive cells in *T. adhaerens*. Of note, DAPI staining was recorded in a fourth channel (not shown), and the unspecific labeling common to all four channels, which corresponds to numerous round-shaped autofluorescent "concrement vacuoles", was subtracted from the image for clarity. (C) Schematic representation of the respective positions of peptidergicimmunoreactive cell populations in *T. adhaerens*, on top and side views, similar to the planes in A. Of note, SITFamide sera also faintly labeled the mitochondrial clusters of fiber cells (not shown). *, upper edge of the animal; le, lower epithelium; ue, upper epithelium. Scale bars: in A, 10 µm for the left and middle panels, 4 µm for the right panel; in B, 10 µm.

Earlier studies using antibodies against RF [15] or FMRF [8] have identified putative peptidergic cells at the rim, and ciliated, endocrine-like cells with neurosecretory features (e.g. expressing SNARE proteins) have also previously been described there and named gland cells [6–8]. Since no FMRFamide peptide precursor could be identified in the *T. adhaerens* genome, we asked whether labelling with anti-FMRFamide antibodies could derive from cross-reactivity with (some of) the populations of newly-identified cells. Upon co-immunolabelling using commercial anti-FMRF antibodies, we observed indeed that SITFamide-expressing cells and, more faintly, TrIns3-expressing cells are also labeled by these antibodies, suggesting that anti-FMRF antibodies actually detect SITFamide (xFigure S1). The remaining three populations of neuropeptide-secreting cells, expressing RWamide, FFNPamide and YYamide distributed away from the rim in more central parts of the organism. Labelling for RWamide is confined to the inside of the organism and does not reach to the surface. The fluorescent peptidergic granules distribute tightly around a rather small nucleus located deep in the middle of the organism. Since they are not observed in correlation with autofluorescent particles nor with large nuclei or cell-bodies, we suggest that RWamide-secreting cells do not correspond to the large and complex fiber cells, landmark of the intermediate, non-epithelial layer [7,8] but to another, as yet unidentified cell type. RWamide-cells spread within a very large ring, sometimes even to the center of the organism (Figure 1C). Finally, cells expressing FFNPamide and YYamide mix and mingle in the center part of the lower epithelium. FFNPamide cells are thin and tall and more abundant than YYamide cells, while the latter appear larger and filled with numerous labelled granules. In this region, cylindrical epithelial cells, lipophil cells and another population of putative endocrine cells (labelled with SNAREs but not FMRFamide antibodies) have been reported [7,8]. Whether FFNPamide and/or YYamide-cells may correspond to SNARE-expressing cells cannot be addressed currently since available antibodies did not allow for double-labelling. *T. adhaerens’s* genome encodes in fact for several isoforms of secretory SNARE proteins [16], and the specificity of antibodies used so far is unclear; it is likely that different populations of secretory cells with different sets of secretory SNARE protein isoforms exist in *T. adhaerens*.

In summary, we uncovered an unexpected diversity of peptidergic cells with unique concentric distribution patterns (Figure 1C) in this morphologically simple animal, indicative of their unique signalling functions potentially regulating many aspects of *T. adhaerens* physiology.

### Neuropeptides elicit strong stereotypical behaviours in *Trichoplax adhaerens*

Placozoans move slowly across surfaces using a densely ciliated lower epithelium [9]. They produce stereotypical behaviours as they flip, fold, flatten, rotate and constantly change their shape [17–20] in a highly coordinated manner, and modulate their speed as a function of light intensity [19] or presence of food [8,18,19,21–23]. Their extra-organismal feeding behaviour is particularly complex [18,21]. How *T. adhaerens* coordinates such movements is only beginning to emerge.

To test the effects of peptides on *T. adhaerens’* behaviour, we recorded over 50 min the motor behaviour of several individuals before and after the application (at min 15) of each one of eleven synthetic neuropeptides predicted from *T. adhaerens* proneuropeptides (ELPE, FFNPamide, LF, LFNE, MFPF, PWN, RWamide, SIFGamide, SITFamide, WPPF, YYamide; Tables S1 & S2; note that the peptides ELPE and MFPF have not been reported so far) at varying concentrations across trials (200 nM - 50 µM; Table S1; in-depth analysis was performed for 20 µM peptide). We observed strong effects on behaviour following peptide treatment, indicating that the peptides reached their targets, as expected from the anatomy of the organism [22]. Several parameters including the area and shape (roundness) of the organisms, as well as locomotion speed and path trajectories were scored. The timing and extent of individual responses varied considerably, making it impractical to reflect the effects in averaged datasets. Hence we show averaged and individual responses as appropriate depending on the nature of the effect of each peptide.

In the majority of cases, the neuropeptides elicited strong changes in movement that were all unique but which we categorized under three general types: (i) crinkling, (ii) rounding up and rotating, and (iii) flattening and churning.

Upon application of PWN, *T. adhaerens* immediately crinkled, as reflected by a sharp decrease in the area of the animals. The effect lasted for approximately 10 minutes, after which the animals recovered their normal shape (Figure 2A, C–D; Movie S1). Application of SIFGamide induced a slower-onset but even stronger crinkling that led to an approximate 50 % decrease in the area, an effect from which the animals only started to recover by the end of the recording. In addition, SIFGamide-treated animals often partially detached from the substrate (Figure 2E, G–H; Movie S2). Detachment was briefly reported elsewhere [24]. These effects are reminiscent of the shape *T. adhaerens* adopts in response to UV light [25] or when flushed with a strong water stream.

**Figure 2.**
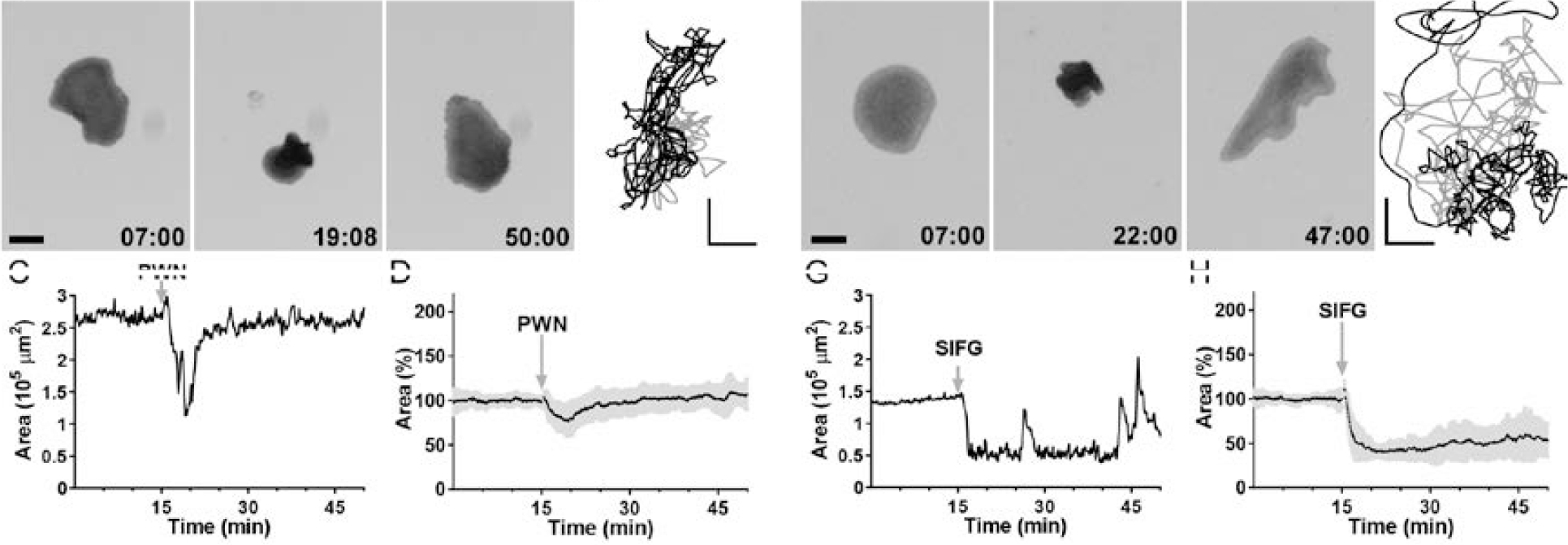
PWN and SIFGamide elicit partial detachment and folding of *Trichoplax adhaerens*. Representative images (A,E), trajectory (B,F) and surface area over time (C,G) of *T. adhaerens* individuals upon a single batch application of 20 µM PWN (A-D) or SIFGamide (E-H). (D,H) Average surface area variations (Mean+StDev) upon 20 µM PWN (n=22 animals in 5 trials) or SIFGamide (n=23 animals in 6 trials). Travelling paths (in grey/black: before/after application) do not show major alteration in trajectories, illustrating that *T. adhaerens* remain attached at the bottom of the dish. PWN acts more transiently than SIFGamide does. Scale bars: 250 µm in A, 200 µm in B and F, 170 µm in E.

The application of LF and LFNE peptides triggered a different type of morphological and behavioural change. Both peptides induced the animals to rotate for several minutes around a fixed axis (arrows in Figure 3E) or to move in circular trajectories (Figure 3B, F; Movies S3 & S4). They transiently flattened, while keeping their round shape, to reach up to 240 % (LF; Figure 3D) or 130 % (LFNE; Figure 3H) of their average area. Upon LF application, animals rapidly adopted a smooth round shape (Figure 3A) and rotated onto them and in the dish. While the animals recovered, their edges were seen undulating at times. Upon LFNE application, the majority of animals rounded up, often adopting a spiral shape (Figure 3E) and turned around themselves (Figure 3F), while a few individuals elongated or undulated at the edges.

**Figure 3.**
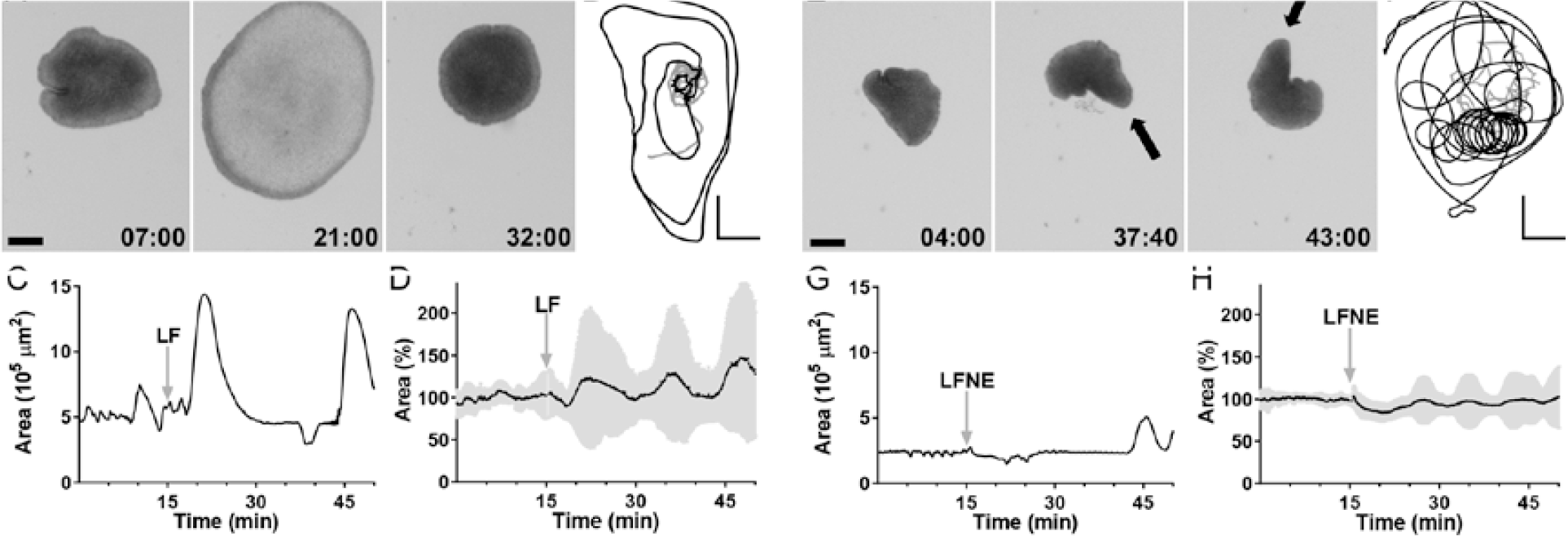
LF and LFNE induce rotation and extreme flattening of *Trichoplax adhaerens*. Representative images (A,E), trajectory (B,F) and surface area over time (C,G) of *T. adhaerens* individuals upon a single batch application of 20 µM LF (A-D) or LFNE (E-H). (D,H) Average surface area variations (Mean + StDev), upon 20 µM LF (n=12 animals in 3 trials) or LFNE (n=22 animals in 5 trials). Travelling paths (in grey/black: before/after application) show drastic changes in trajectories upon peptide application. Not only are individuals rotating around a fixed axis in an unusually "frozen" shape (see arrows in E), but also moving in small circular paths within the dish. They periodically underwent massive flattening. Scale bars: 250 µm in A and E, 200 µm in B and F.

Another group of peptides, comprising FFNPamide, ELPE, MFPF and WPPF, induced actions reminiscent of feeding behaviour, without entirely recapitulating the feeding sequence (Movies S5-8). When feeding, *T. adhaerens* stops moving before a large fringe of its periphery stretches and flattens down, adhering to the bottom, while its central part undergoes coordinated rhythmic movements including stretching, contracting and rotating components ("churning" [18,21]), then loosens itself from the bottom and resumes crawling. All four peptides increased the number of flattening events (Figure 4C, G, K, O), as shown by an increase in average area (Figure S2). Most often, flattening and pausing events occurred concomitantly (Figure 4B vs. D, J vs. L, N vs. P), while they were occasionally slightly shifted (Figure 4F vs. H). Flattening induced by FFNP was extreme, as the centre part of the organism appeared extremely pale, suggesting that the animal is able to thin out to an extent that has not been reported before - up to 5 times its "default" size. ELPE, MFPF and WPPF induced the animals to adhere more tightly to the dish at their rim (which hence appears paler (Figure 4E, I, M)), to flatten down and to churn with slightly different movements. In the presence of MFPF, animals churn slower or faster than untreated animals, and for extended periods of time. FFNPamide addition also triggered moderate churning movements. Interestingly, the numerous FFNPamideimmunoreactive cells embedded in the lower epithelium (Figure 1) are found in the same area as the digestive enzyme-releasing lipophil cells [8,21], hence are well positioned to fulfil a role in modulating feeding behaviour [4,26]. Altogether, these behaviours could be subprograms of the motor alterations associated with the food uptake process; whether the peptides involved are acting independently, sequentially or in synergy requires further investigation.

**Figure 4.**
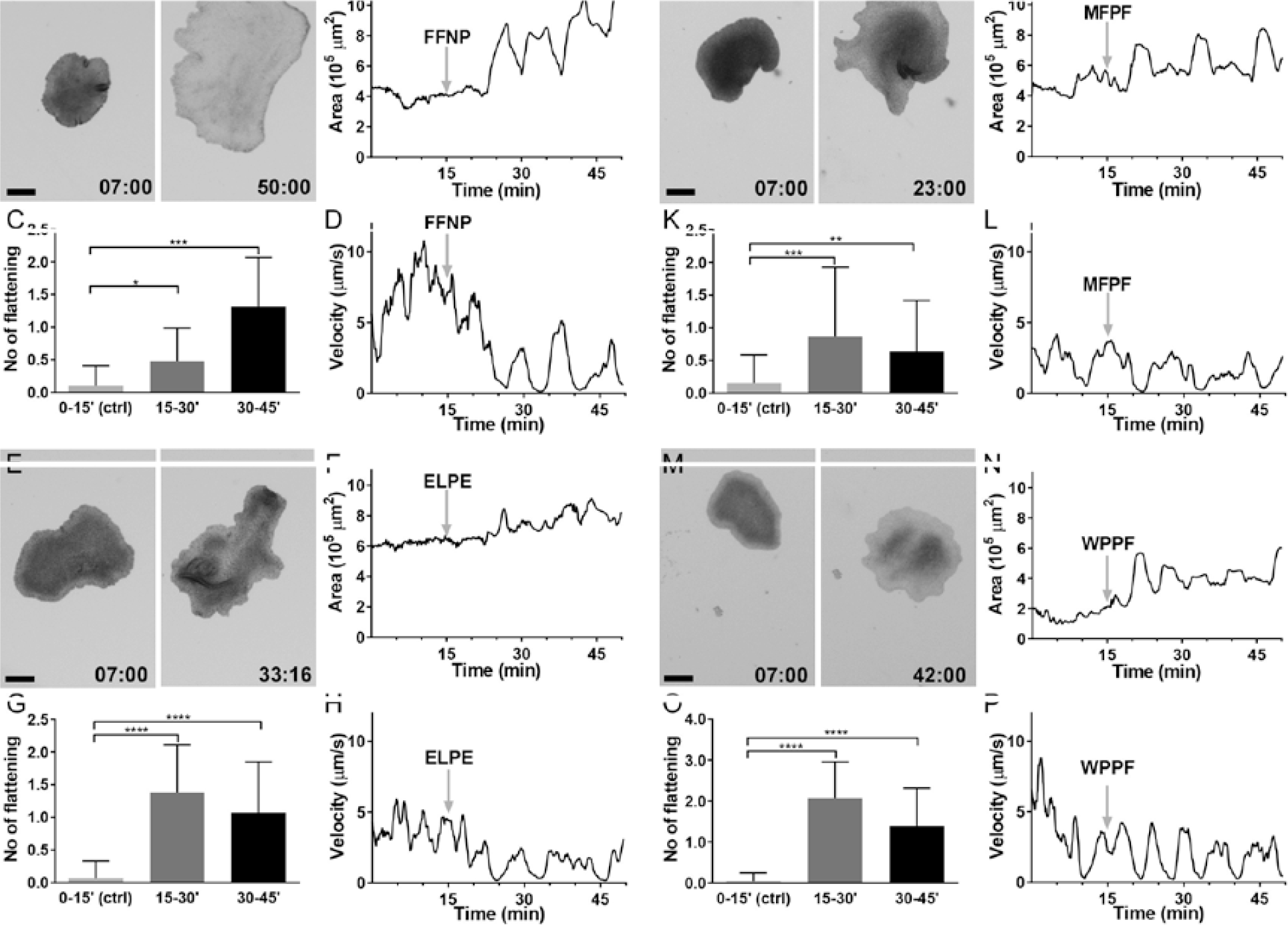
FFNPamide, ELPE, MFPF and WPPF provoke flattening and different forms of "internal" movement, e.g. churning, in *Trichoplax adhaerens*. Responses of *T. adhaerens* to a single application of 20 µM FFNPamide (A-D), ELPE (E-H), MFPF (I-L) or WPPF (M-P). Representative images (A,E,I,M), area (B,F,J,N) and speed (D,H,L,P) over time of an example individual. Average numbers of flattening events (C,G,K,O) before (0-15’(ctrl)) and after peptide application (15-30’, 30-45’)(Mean+StDev). For FFNPamide (C), n=20 animals in 3 trials; for ELPE (G), n=42 animals in 5 trials; for MFPF (K), n=39 animals in 5 trials; for WPPF (O), n=46 animals in 4 trials. While all 4 peptides invariably yield more frequent flattening in *T. adhaerens*, each of them induces specific alterations in shape and movements that can be observed in Suppl. video material. Scale bars: 250 µm in A, E and I, 170 µm in M. Statistical tests applied: Wilcoxon for FFNPamide, ELPE, WPPF; Mann-Whitney for MFPF; ns, non significant; *, p<0.05; **, p<0.01; ***, p<0.001; ****, p<0.0001.

Finally, a few peptides induced only subtle alterations in *T. adhaerens* behaviour (Figure S3; Movies S9-11). Neither SITFamide, YY, nor different RWamide peptides affected the flattening rate of the animals. SITFamide induced an effect which was difficult to quantify. The speed of the animals was significantly lower during the 30 minutes monitored after SITFamide application. Some animals slowed down (Figure S3D) without stopping (Figure S3A, B). In one case, an animal stopped but did not flatten (2_102, Figure S3). Application of YYamide resulted in a small but significant increase in the speed of or the distance travelled by the animals during the later phase of the recordings (Figure S2G). In one case, an animal increased its area without slowing down (1_128, Figure S3). Finally, the effects of RWamides were tested in a pilot series of experiments using the three slightly different peptides RWamide1, RWamide2, RWamide3 and their respective forms bearing a N-terminal pyroGlutamine modification pGluRWamide1, pGluRWamide2 and pGluRWamide3. No drastic effect was observed over the 30 minutes of recordings. Hence, pRWamide2 was arbitrarily selected for further trials. No change in area, trajectory, speed or distance travelled was noted. Application of the vehicle alone did not alter the animals’ behaviour (Figure S4; Movies S12; S13).

Overall, our data show that most predicted neuropeptides are able to strongly alter *T. adhaerens* behaviour, suggesting that peptidergic signalling pathways that can affect the animals’ contractility and movements are in place. Moreover, we describe a previously undocumented capacity of extreme physical alterations of *T. adhaerens*, from extreme flattening to crinkling and from churning to rotating. Our results suggest that motor behaviours in *T. adhaerens* likely result from the coordinated interplay of several cellular arrays that are under the strong influence of peptidergic signals.

An earlier study reports that gland cells secrete an endomorphin-like peptide, which arrest ciliary beating, inducing feeding-like pauses [24] (Figure S5). Interestingly, we observed that the application of the endomorphin-like peptide also induced flattening but no churning (Table S1). At lower concentration, crinkling was observed, whereas high concentrations led the animal to rotate. While it is conceivable that some of the changes in shape and movements reported herein may rely in part on an arrest ciliary beating, they might also be generated by fast contractions of epithelial cells [20] or by contractions of the large, actin/microtubule-rich fiber cells in between the two epithelial layers [26,27], which are connected to each other and can reach out to all cell types [8].

Peptide hormones and their respective receptors appeared early in metazoan evolution and have been described in all animals with a nervous system including ctenophores [13,28] and cnidarians. Our study suggests that peptide hormones play an important role in cell-cell communication in *T. adhaerens* as well. Of note, the genome of *T. adhaerens* encodes for a rich repertoire of G-protein coupled receptors [10,29], at least some of which likely act as neuropeptide receptors [11]. The neuropeptide repertoire in *T. adhaerens* is less complex than that of eumetazoans and might thus represent a more ancestral stage [11]. It will be interesting to explore the conservation pattern of the *T. adhaerens* neuropeptides in other placozoans [30,31]. Since the *T. adhaerens* genome also contains all determinants of transmission via classical amino-acid neurotransmitters [10], it is essential to scrutinize whether the animal combines elements of peptide hormone and classical amino-acid transmission, and whether the peptide-containing cells described herein might represent specialized, non-neuronal secretory cells or neuroendocrine-like/neuron-like cells able to co-release both types of transmitters, as is found to occur in neurons and endocrine cells in animals with nervous systems [32].

Altogether, our data demonstrate that cell type diversity in placozoans is higher than previously inferred [7,8] and that several specialized cell types, regarded as the "basal building blocks of multicellular organisms" [33], have already emerged in this phylum. Indeed, a recent single-cell transcriptome study confirmed the presence of several hitherto unexplored lower-frequency cell types expressing unique signalling peptides [34]. In *T. adhaerens*, several populations of topographically organized peptide-secreting cells can strongly modulate the animals’ movements. Our results suggest that peptidergic signalling is an important mode of communication for placozoans, as is the case for cnidarians and bilaterians [11,35–39].

## EXPERIMENTAL PROCEDURES

### Animal Maintenance

*Trichoplax adhaerens* (Grell strain, haplotype H1) were maintained in large glass Petri dishes (15-cm diameter) containing artificial seawater (filtered ASW, salinity of 1025 ppm (38 g/l) salinity; Coral Pro Salt, Red Sea), at 26°C and under a 12:12 hr constant light cycle. They were fed once a week with 8-10 ml of a 1:1 mixture of *Nannochloropsis* (Florida Aqua Farms) and *Pyrenomonas helgolandii* (SAG, Göttingen, Germany) cultured separately in ASW complemented with 0.03 % of Micro Algae Grow (Florida Aqua Farms). About 80 % of the medium was replaced every other week. When reaching a population density of over 100-200 individuals per dish, 10 to 20 animals were randomly taken up with a pipette to set up a new dish.

### Antibody production

Polyclonal antibodies were generated in rabbits (2 rabbits per peptide, Speedy program, Eurogentec) against peptide candidates [11,12] that were predicted to be amidated, namely "FFNPamide" (CQFFNP-amide), "RWamide" (CRDQPPRW-amide), "SIFGamide" (CQANLKSIFG-amide), "SITFamide" (CNSESTQQGIPSITF-amide), "YYamide" (CGYDDYYY-amide), and Insulin-3 "TrIns3" (CPIH-amide); for immunization, peptides were coupled to ovalbumin as carrier protein; the cysteine residue added at the N-terminal of the peptides allows for subsequent immobilization on a Sulfolink resin (ThermoFisher). Sera were affinity-purified as described [14], each resulting in two fractions after elution at pH2.9 and pH2.3. Initial immunostainings were carried out with crude and purified antibodies. The two sera raised against each peptide gave similar staining patterns; In all cases, stainings were invariably abolished upon preincubation of the primary antibody with the peptide it was raised against.

### Immunolabelling

*T. adhaerens* were left to adhere at the bottom of 35-mm plastic Petri dish filled with ASW, swiftly replaced by 4 % paraformaldehyde (PFA) with Dextran in ASW for fixation (1-2 hrs, RT). After brief washes in ASW followed by 0.1 M phosphate buffer pH7.4 (PB) and a 1 hr-blocking step in 3 % normal goat serum (NGS) + 0.2 % Triton X-100 in PB, they were incubated for 2 hrs in primary antibody (anti-peptide: 1 µg/ml; anti-FMRF (1:300; Enzo Life Sciences #BML-FA1155 or Phoenix Pharmaceuticals #H-047-29) in blocking solution. After thorough washes in PB, a 1-hr incubation in the corresponding secondary antibody coupled to a fluorophore (1:600, GAR-Alexa Fluor, Molecular Probes) and further washes in PB, they were incubated 2 minutes in Hoechst (1:10,000; Molecular Probes), rinsed and mounted with Prolong-Gold (LifeTech) on glass slides. For co-labelling with antibodies raised in the same species, *T. adhaerens* were instead incubated 45 minutes in a mix of antibodies coupled to Alexa fluorophores using the Zenon labelling kit (LifeTechnologies) following manufacturer’s instructions, washed in PB, post-fixed 15 minutes in 4 % PFA, washed again and incubated in Hoechst before being mounted on a slide. For blocking experiments, we preincubated the antibodies in three times excess of the respective peptides for 2 h before immunostainings.

### Imaging

Specimens were imaged on an inverted confocal microscope set-up (Zeiss LSM 780) using a 63x-oil immersion objective (NA1.4). Images stacks were acquired over the entire thickness of the animal with optical slices of 1.3 µm with an overlap of 0.75 µm, with each fluorophore imaged separately.

### Behavioural testing

Most peptides were dissolved in water, except for YYamide and LF, which were dissolved in 5 mM NH_4_HCO_3_. 10-15 *T. adhaerens* were pipetted into a 35-mm plastic Petri dish filled with 3 ml ASW and allowed to settle for 15-30 minutes at 26°C. The dish was then transferred to the stage of a stereomicroscope (Nikon SM225 coupled to a Nikon DSR12 camera) with a zoom of 0.63x or 3x, and imaged for 50 minutes. After 15 minutes, each peptide (200 nM - 50 µM from a 10 mM stock solution) was applied once and swiftly mixed in the bath by gentle pipetting. Peptides tested were the following: "ELPE" (GKSFELPE), "FFNPamide" (DDQFFNP-amide), Insulin-3 "TrIns3" (CPIH-amide), "LF" (DDSQDGYALF), "LFNE" (QEPGISLFNE), "MFPF" (EDDLPGMFPF), "PWN" (EQGALLDIPWN), "RWamide1" (DQPPRW-amide), "RWamide2" (DQPTRW-amide), "RWamide3" (DQPSRW-amide), RWamides bearing a N-term pyroGlutamine modification "pGluRWamide1-3", "SIFGamide" (EDQANLKSIFG-amide), "SITFamide" (NSESTQQGIPSITF-amide), "WPPF" (EDQQNKPYNGWPPF), and "YYamide" (DYDDYYY-amide) (see Table S1 and S2 for details). Control experiments were carried out by application of the vehicles only. Endomorphin-2 (Genscript, RP10926) was tested at concentrations of 1, 15 and 50 µM. Images were taken every 4 seconds under moderate and constant illumination, and at a room temperature of approximately 20°C. Upon recording, the dishes were placed back in the incubator with food to check animal viability 24 hrs after treatment.

### Image Analysis and Statistical Methods

Black and white image series (Tif) were analysed with Fiji. Only individuals that were visible during the entire time of the recording were tracked. Since animals in close contact may coordinate their behaviours [17,24], only animals which were not in contact with one another were monitored to avoid oversampling. In rare cases when individuals were in contact with each other, only one was monitored. Images on which the application pipettes appeared were not taken into account for analysis. The maximum trajectory area was set as ROI for each individual and a time stamp applied (see .avi videos as Supplemental information). A threshold (Yen, Fiji) was set to optimally define the contour of each individual, allowing for the binarization of the image and measurement of the animal’s size (area) using the Plugin ‘Analyse Particles’ in Fiji. Trajectory, distance covered and speed were assessed from the centroid, using the ‘Mtrack2’ Plugin in Fiji. To standardize area variations over several animals, the average area was defined for each individual over the first 15 minutes of recording and used as a baseline to normalize the values over the recording. Standard deviations /Standard Errors of the Mean were calculated for each time point for all individuals using Excel or Prism. Results are presented as Mean ± StDev in Figure 4, Mean ± SEM in Figures S3 and S5. When the values followed a normal distribution, a paired t-test was used (Figures S3 and S5). Otherwise, a paired Wilcoxon test was applied (Figure 4).

## AUTHOR CONTRIBUTIONS

FV, EW, GJ, DF designed the study; FV, EW, LT, SG performed experiments; KK, BS provided strain and advice; FV, EW, LT, SG, BS, GJ, DF analysed data; FV, EW, GJ, DF wrote the paper and all authors reviewed, commented on, and edited the manuscript.

## ACKNOWLEDGMENTS

This work was supported by the Swiss National Science Foundation (Marie-Heim Vögtlin grant PMPDP3_158377 to FV and grant 31003A_160343 to DF). EAW was supported by a grant from the DFG - Deutsche Forschungsgemeinschaft (Reference no. JE 777/1). We thank Yves Mingard, Dominique Trauffer, and Marie-Laure Gadolini for taking care of our *Trichoplax* lines and technical assistance, and Mathieu Künzi for support with analysis and reading of the manuscript.

## SUPPLEMENTAL INFORMATION

**Figure S1.**
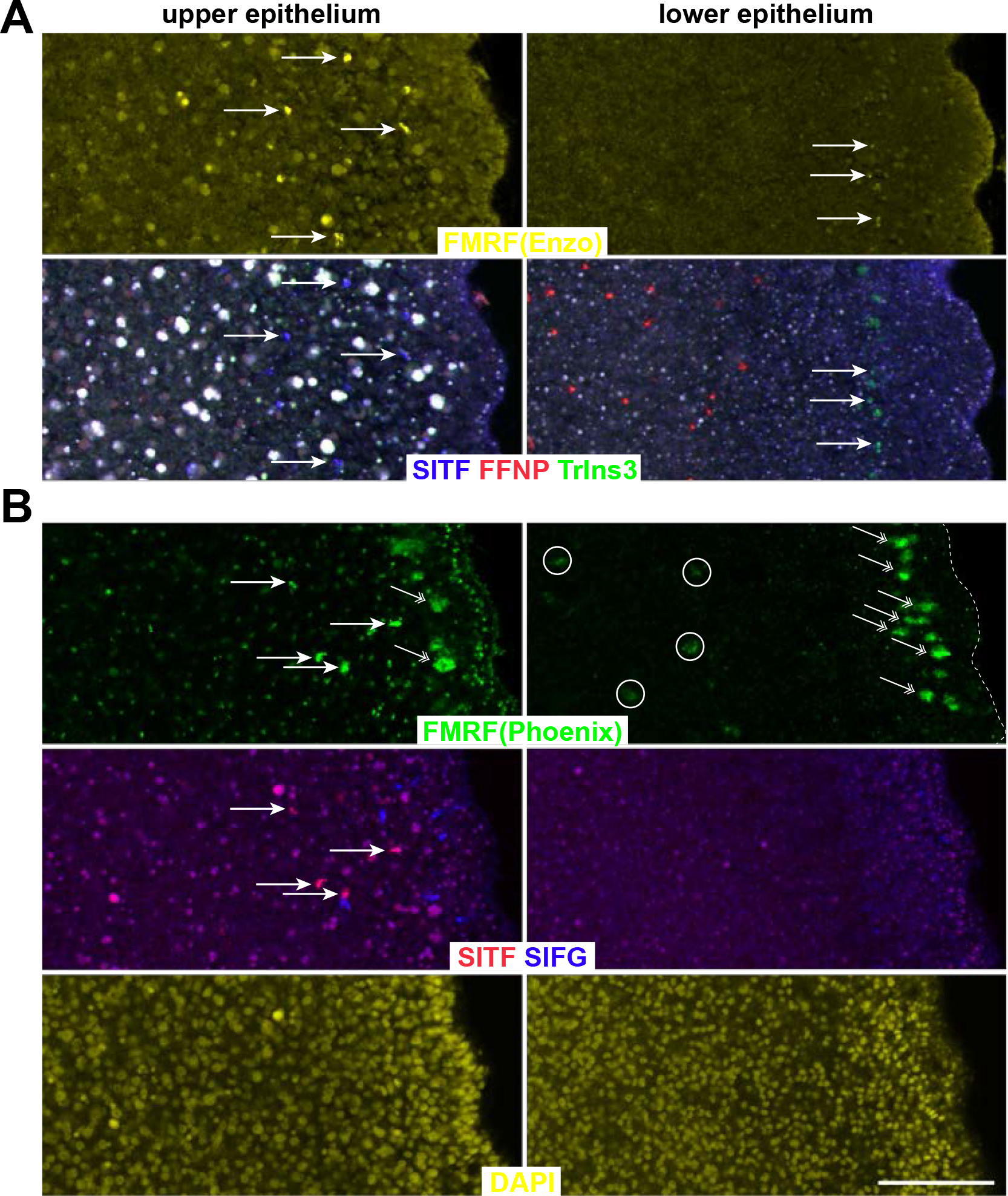
Different antibodies against FMRFamide co-label SITFamide-immunoreactive cells. Anti-FMRFamide antibodies from Enzo (#BML-FA1155; yellow in A) and from Phoenix labs (#H-047-29, lot 01479-1; green in B) were used in combination with other peptide antibodies. Labeling is illustrated in a single optical slice of the upper epithelium (left column) and the lower epithelium (right column) of a same image stack. (A) Quadruple staining for FMRFamide from Enzo (FMRF(Enzo), yellow, upper panels), SITFamide, FFNPamide and TrIns3 (blue, red and green, lower panels) shows that this FMRFantibody strongly co-labels SITFamide-immunoreactive cells and to a lesser extent, TrIns3-stained cells (arrows). Populations of immunoreactive cells for SITFamide, FFNPamide and TrIns3 do not overlap. Of note, the numerous round-shaped white particles observed predominantly in the upper epithelium are autofluorescent "concrement vacuoles". (B) Quadruple staining for FMRFamide from Phoenix (FMRF(Phoenix), green, upper panels), SITFamide and SIFGamide (red and blue, middle panels) and Hoechst (yellow, lower panels) shows that this FMRF antibody also strongly co-labels SITFamide-immunoreactive cells (arrows) but not SIFGamide-stained cells. This antibody also labels a row of large cells located close to the edge of the organism (double-arrowheads) and some scarce cells of the lower epithelium (circles) likely corresponding to another population of unidentified gland cells [8], as well as small unidentified elements in the upper epithelium. Populations of immunoreactive cells for SITFamide and SIFG do not overlap. Note again the presence of autofluorescent particles, visible in all channels, and the difference of Hoechst-stained nuclei size and number across both epithelia. Scale bar: 30 µm.

**Figure S2.**
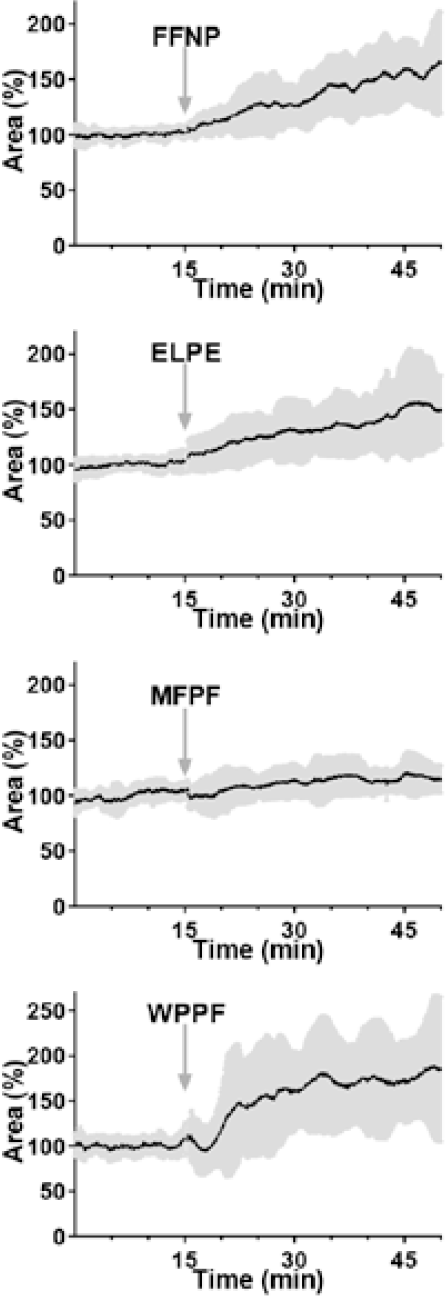
Average area over time upon application of FFNPamide (n=17), ELPE (n=24), MFPF (n=17), WPPF (n=18). A single peptide application (arrow) induced a sustained increase of the standard deviation, reflecting an effect of the peptide.

**Figure S3.**
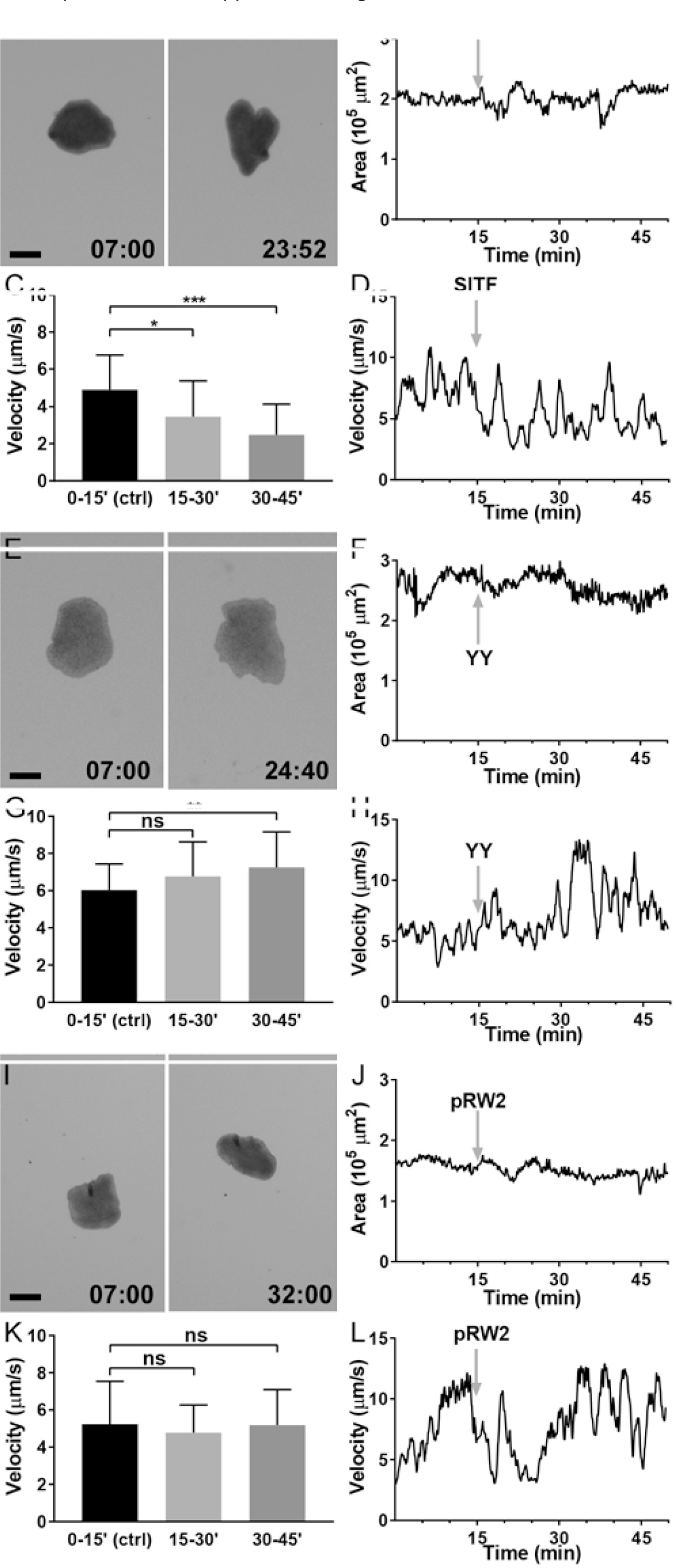
Subtle effects of SITFamide, YYamide or pRWamide2 application to *Trichoplax adhaerens*. Responses of *T. adhaerens* to a single application of 20 µM SITFamide (A-D), YYamide (E-H) or pRWamide2 (I-L). Representative images (A,E,I), area (B,F,J) and speed (D,H,L) over time of an example individual. (C,G,K) Average velocities (C, SITFamide, n=15; G, YYamide, n=15; pRWamide2, n=15) before (0-15’(ctrl)) and after peptide application (15-30’, 30-45’) (Mean+StDev). Scale bars: 250 µm in A, E and I. Statistical tests applied: Wilcoxon for SITFamide, Mann-Whitney for YYamide and pRWamide2; ns, not significant; *, p<0.05; **, p<0.01; ***, p<0.001; ****, p<0.0001.

**Figure S4.**
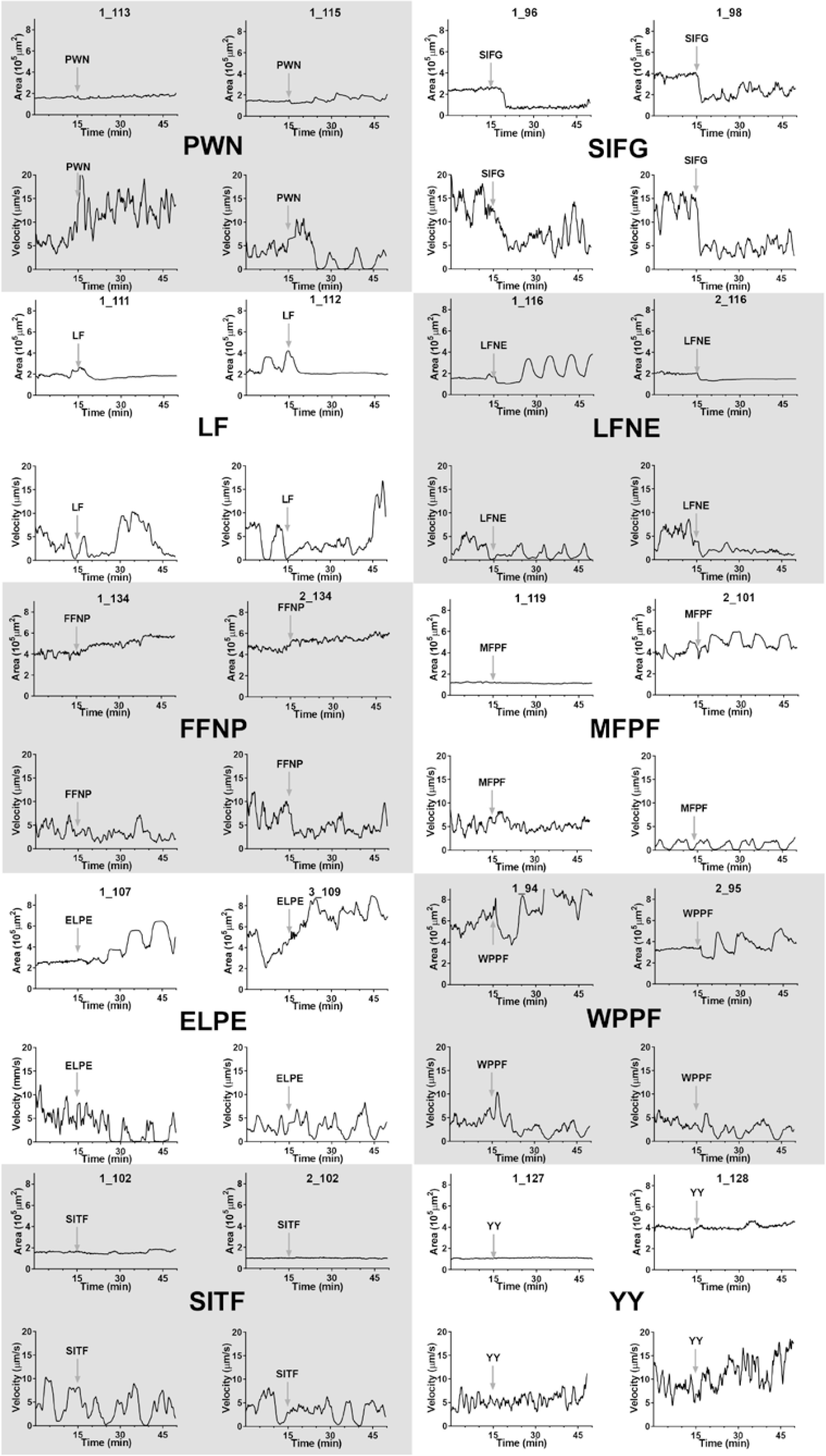
Example traces of behaviours (area and speed) of *T. adhaerens* individuals upon application of the indicated peptides.

**Figure S5.**
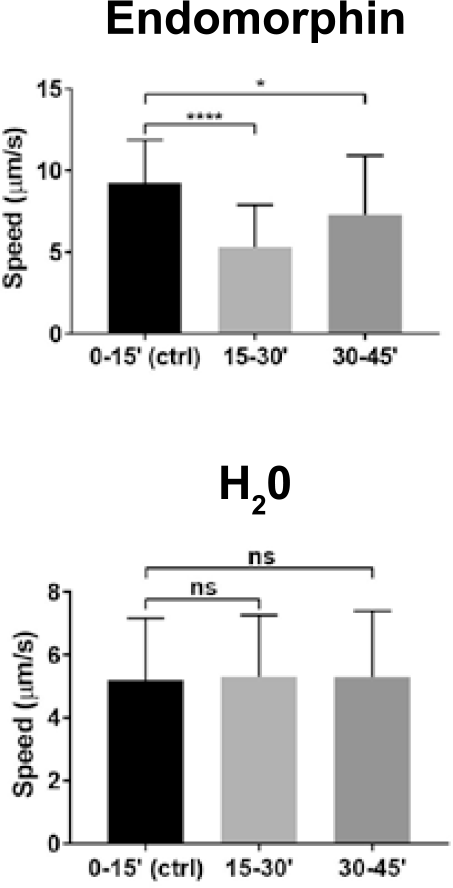
Average speed upon application of endomorphin-2 (n=14) or H_2_O (n=16). Average speed before (0-15’(ctrl)) and after H_2_O or endomorphin application (15-30’, 30-45’) (Mean+SEM). Statistical test applied: Paired t-test; ns, not significant; *, p<0.05; **, p<0.01; ***, p<0.001; ****, p<0.0001.

**Table S1.**
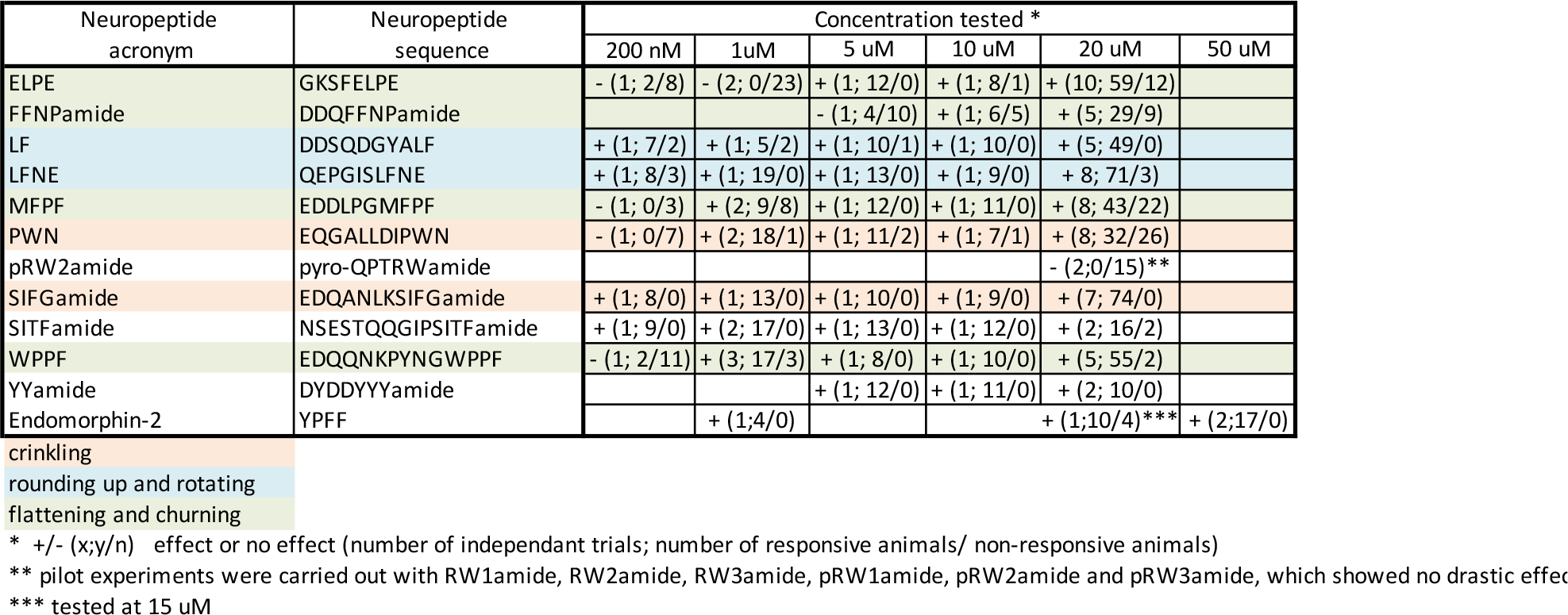
Summary of the neuropeptides tested. The number of independent trials carried out for a given neuropeptide at a given concentration, as well as the number of animals showing or not the described phenotype (see color code) are given. Note that the numbers of individuals reported in the main analyses were often lower as only animals observable during the entire time of the recording and not touching each other were used for precise quantification.

**Table S2.**
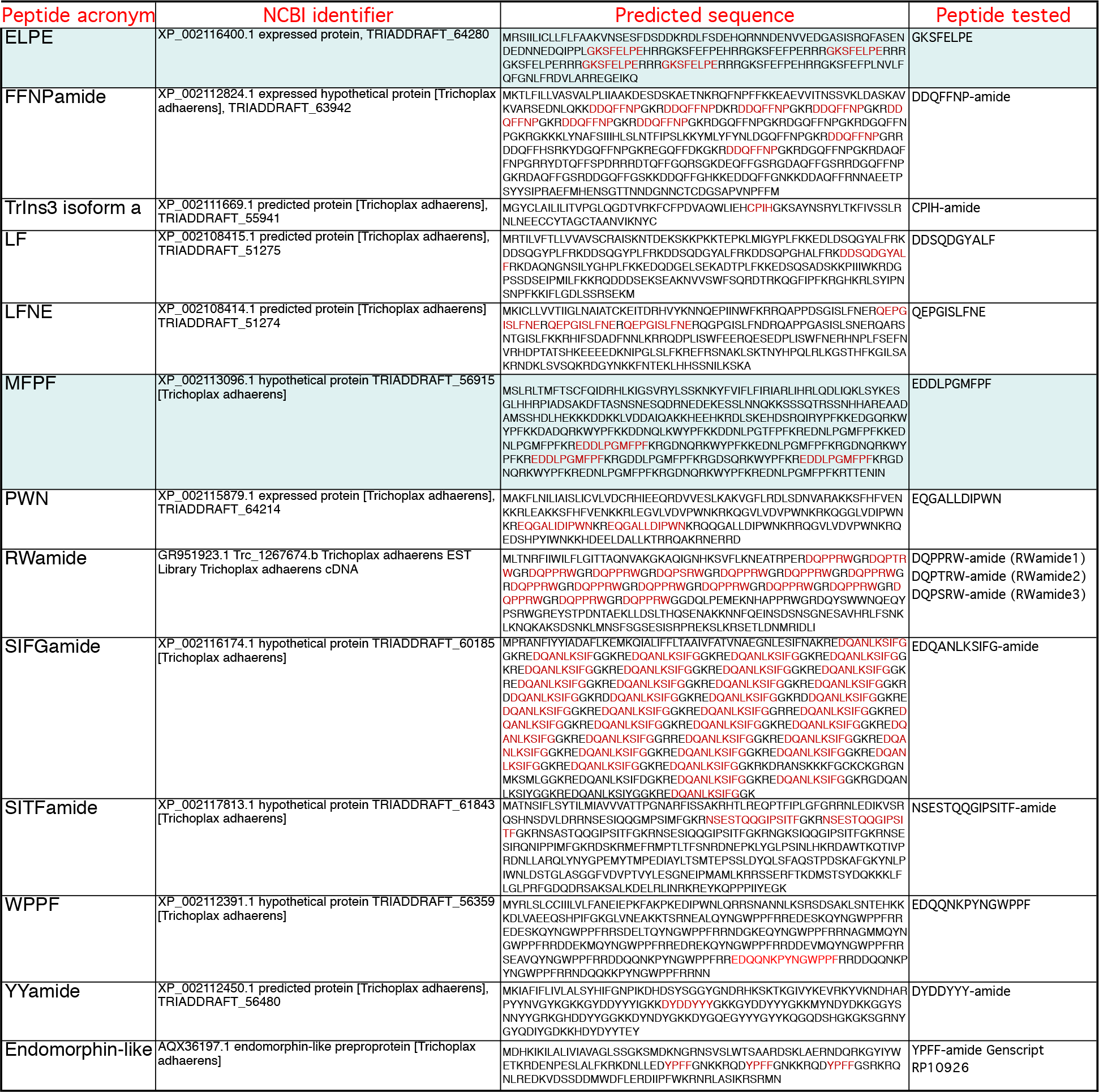
List of neuropeptides precursor sequences. Two of the neuropeptides tested are novel and highlighted in blue. The epitopes used for generating antibodies are highlighted in red in the predicted sequence.

**Movie S1**: Time-stamped videos showing examples of *T. adhaerens*’s behaviour upon application of 20 µM PWN.

**Movie S2**: Time-stamped videos showing examples of *T. adhaerens*’s behaviour upon application of 20 µM SIFGamide.

**Movie S3**: Time-stamped videos showing examples of *T. adhaerens*’s behaviour upon application of 20 µM LF.

**Movie S4**: Time-stamped videos showing examples of *T. adhaerens*’s behaviour upon application of 20 µM LFNE.

**Movie S5**: Time-stamped videos showing examples of *T. adhaerens*’s behaviour upon application of 20 µM FFNPamide.

**Movie S6**: Time-stamped videos showing examples of *T. adhaerens*’s behaviour upon application of 20 µM ELPE.

**Movie S7**: Time-stamped videos showing examples of *T. adhaerens*’s behaviour upon application of 20 µM MFPF.

**Movie S8**: Time-stamped videos showing examples of *T. adhaerens*’s behaviour upon application of 20 µM WPPF.

**Movie S9**: Time-stamped videos showing examples of *T. adhaerens*’s behaviour upon application of 20 µM SITFamide.

**Movie S10**: Time-stamped videos showing examples of *T. adhaerens*’s behaviour upon application of 20 µM YYamide.

**Movie S11**: Time-stamped videos showing examples of *T. adhaerens*’s behaviour upon application of 20 µM pRWamide2.

**Movie S12**: Time-stamped videos showing examples of *T. adhaerens*’s behaviour upon application of H_2_O.

**Movie S13**: Time-stamped videos showing examples of *T. adhaerens*’s behaviour upon application of NH_4_HCO_3_.

